# Multi-Unit Activity contains information about spatial stimulus structure in mouse primary visual cortex

**DOI:** 10.1101/029108

**Authors:** Marie Tolkiehn, Simon R. Schultz

## Abstract

This study investigates the spatial and directional tuning of Multi-Unit Activity (MUA) in mouse primary visual cortex and how MUA can reflect spatiotemporal structures contained in moving gratings. Analysis of multi-shank laminar electrophysiological recordings from mouse primary visual cortex indicates a directional preference for moving gratings around 180°, while preferred spatial frequency peaks around 0.02 cycles per degree, which is similar as reported in single-unit studies. Using only features from MUA, we further achieved a significant performance in decoding spatial frequency or direction of moving gratings, with average decoding performances of up to 58.54% for 8 directions, and 44% correctly identified spatial frequencies against chance level of 16.7%.

## I. Introduction

The cortical microcircuit governs how we see, hear and think. Reverse-engineering the functionality of this circuit is a major project of modern neuroscience, and of the emerging field of neural engineering. The mouse primary visual cortex is a prime candidate for studying the principles of information processing in a cortical circuit, as it possesses a similar range of cell types and receptive field classes to that of other mammals [1], while allowing numerous recently developed molecular and optogenetic tools to be applied. In order to understand how the cortical circuit functions, and changes during learning and memory processes, it is essential to analyse its operation during behaviour, and in particular by recording how activity in different elements of the circuit changes over behavioural session. Unfortunately, the standard technique for observing neural activity, single unit neurophysiological recording (single unit activity, SUA), is not particularly well suited to chronic recording strategies spanning multiple behavioural sessions, as individual units are lost due to drift or damage if recording probes are left *in situ* [2], [3].

Alternatives to single unit recording include thresholding the signal on each channel of a multi-electrode recording array, to collect Multi Unit Activity (MUA), and collecting low frequency electrical signals from each channel (local field potentials, or LFP). Both of these measures integrate signals across a small volume of tissue, and show improved robustness in comparison to SUA over multiple recording sessions. This has led to their use in Brain-Machine Interface (BMI) applications [4], [5], and suggests that they may be useful probing the neural substrates of learning mechanisms in the cortical circuit.

One drawback is that mouse visual cortex, unlike that of cats and primates, does not show orientation columnar organisation, but instead exhibits a salt-and-pepper organisation [6], [7], whilst still displaying a high orientation selectivity [8], [1], [7]. Because of this fine-scale random organisation, it might be expected that MUA and LFP in mouse visual cortex contain minimal information about the spatial structure of a stimulus beyond retinotopy. However, we conjecture that a residual bias in the orientation or spatial frequency tuning sampled by the MUA or LFP on a single channel may leave a substantial amount of information. Indeed, it has recently been shown in mouse hippocampus (which also shows a salt-and-pepper organisation) that LFP can be accurately used to decode spatial position [9].

In this paper, we demonstrate that individual MUA channels contain a significant amount of information about the direction and spatial frequency of a drifting grating. We show that the pattern of MUA across channels can be used to decode direction and spatial frequency with performance substantially above chance, and conjecture that this may provide an extremely useful tool for probing changes in cortical circuit information representation during behavioural learning paradigms.

## II. Methods

### A. Electrophysiology

10 acute electrophysiological experiments were performed on female, young adult (age 2-4 months) wild-type C57Bl/6 mice (Harlan, UK) with a target area of left primary visual cortex. The animals were used in accordance with protocols approved by UK Home Office project and personal licenses. The mice were kept in Imperial College animal facility under an inverted 12-hour light/dark cycle. All electrophysiological recordings were carried out during the dark phase of the cycle.

The isoflurane-anaesthetised mouse was placed 25cm away from the monitor (Samsung 2233Z), covering approximately 60°x75° of the visual field. The left eye was treated with eye gel and covered with black tape to avoid confounding effects attributable to the binocular zone of vision or other visual inputs.

The Neuronexus A4x8 silicon microelectrode was lowered slowly into the brain to a depth between 800*μ*m and 1050*μ*m, at a speed of a few 10*μ*m/s. The probe was equipped with four shanks, 200*μ*m apart, with 8 linearly arranged recording sites of 177*μ*m^2^, each separated by 100*μ*m.

### B. Stimuli

In this study, we presented a set of sinusoidal drifting gratings, 20 repetitions each, of 6 different Spatial Frequencies (SF) at 8 different directions, interleaved with 1 minute spontaneous activity (1s stimulus-ON time, with 1s pre-stimulus time), at a constant temporal frequency of 1.6 Hz. SF were given at [0.01, 0.02, 0.04, 0.08, 0.16, 0.32] cycles per degree (cpd), and directions were equidistantly spaced at 45° from 0° to 315°. 0° corresponds to rightward moving, 90^°^ to upward moving, and 180^°^ to leftward moving gratings.

Stimuli were generated with FlyMouse, software refined from FlyFly, a Matlab Psychophysics Toolbox based interface developed by the Motion Vision Group at Uppsala University (http://www.flyfly.se/about.html) and customized by Silvia Ardila Jiminez and Marie Tolkiehn.

### C. Data Acquisition

Signals were acquired by Ripple Grapevine (Scout Processor), amplified with a single-reference amplifier with onboard filtering and digitisation (Grapevine Nano front end), and software Trellis, which enabled for a live display of the channels. Synchronisation between screen output and stimulus presentation was assured by a custom-made photodiode circuit board and photosensor (LCM555CN).

## III. Analysis

*1) Multi-Unit Analysis:* For the Multi-Unit (MU) analysis, we first analysed the visual responsiveness of each electrode site. A one-way ANOVA identified those electrode sites, which showed a significant change in average firing rate between the pre-stimulus time (1s) and during stimulus presentation (1s). This resulted in 67.8% of the channels (217 of 320 electrode sites) exhibiting a significant difference in firing. For further analysis we only kept the sites showing a significant stimulus modulation.

In order to estimate which stimulus elicited a significant response, we performed a two-tailed t-test on the mean ON and OFF firing rates over the 48 stimuli. This resulted in up to 16 stimuli being rejected, which was also evident from raster plots (not shown).

*2) Decoding Feature Selection:* For the multinomial classification of the 48 stimuli, we evaluated decoding performance using a variety of classifiers such as Linear Discriminant Analysis (LDA), Naive Bayes, kd-Tree and k-Nearest neighbour, on two types of features: a) binned spikes at 20 ms and b) spike count. We chose these features over a more complicated feature extraction at this stage as a proof of concept that MU signals can be used to significantly decode different stimuli from these types of neural responses.

The decoding task was comprised of 2 parts for each feature type: *A)* Decoding the Spatial Frequency (1 of 6 SFs), and *B)* Decoding the direction from the responses (1 of 8 directions).

Classification accuracy was evaluated against chance level, validated with a 2-fold cross-validation and averaged over 20 repetitions with random permutations of the partitions. Decoders were tested at 8 directions at the SF of 0.02 cpd, and in the SF decoding regime for 6 SF at 180° to ensure the stimuli were in a detectable range. All stimulus cases had equal probability. This means that chance probability varied between 16.67% for SF to 12.5% for the directions. Consistent classification above chance level suggests the decoder's successful use of inherent structures about SF or direction in the MUA.

*3) Multi-Unit Tuning Properties:* Direction tuning of the MU was evaluated with the sum of two modified von Mises functions similar to those described in [8], [10]. SF tuning curves were fit with two modified log-normal functions. In both cases, these fits were used to estimate peak response and cutoff-frequency. Orientation and direction selectivity were determined with the Orientation Selectivity Index (OSI) and Direction Selectivity Index (DSI), which were defined as *OSI* = (*R_pref_ + R_null_ −* (*R_ortho+_ + R_ortho−_*))/(*R_pref_ + R_null_*), with *R_pref_* as the preferred direction, *R_null_* the opposite direction, and *R_ortho±_* denoting the orthogonal directions [11]. DSI was defined as *DSI =* (*R_pref_ − R_null_)/R_pref_*. An OSI of 1 represents high selectivity, an OSI of 0 means each stimulus produces an equal numbers of spikes.

## IV. Results

Investigating the spike responses to 48 different stimuli showed that most channels were visually responsive. Figure 1 reveals a raster plot of 20 repetitions of one stimulus for one mouse. All 32 channels are shown, where different colours represent different electrode sites. The top red line indicates the period the stimulus (leftward moving grating at 0.02 cpd) was on. It is evident from the raster plot that most channels were consistently visually responsive and that their activity was highly modulated by this stimulus.

**Fig. 1.**
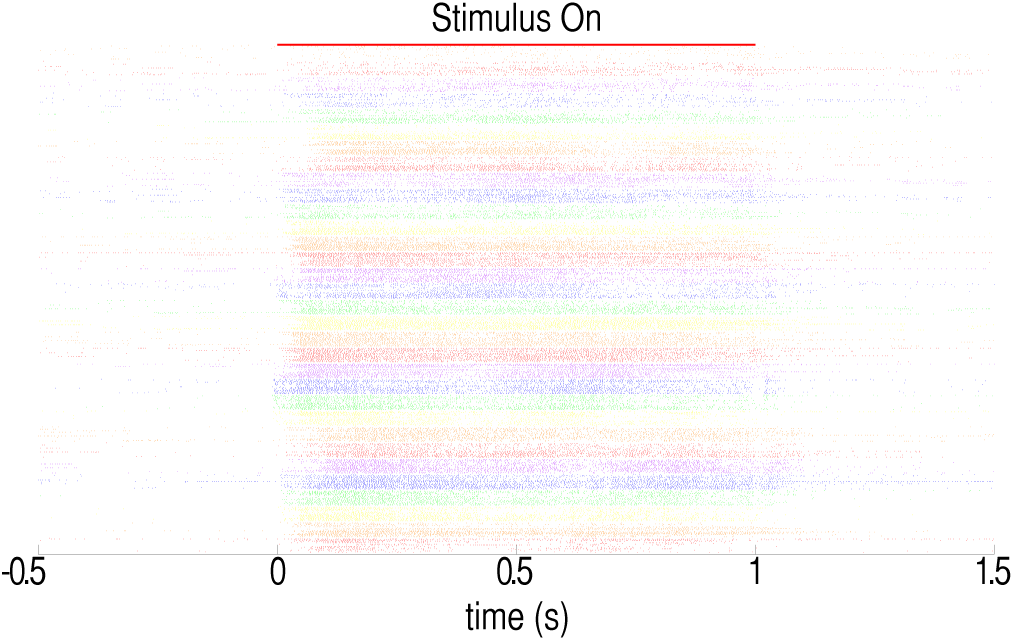
A high fraction of channels is stimulus responsive. Raster plot of MUA of 32 channels to 20 repetitions of a leftward moving grating. Different channels are illustrated by different colours. Each line highlights a spike incident. Top red line top indicates stimulus ON time.

### A. Direction Tuning

A number of MU showed to be significantly modulated by moving gratings. An example of two strongly tuned MU is presented in Fig. 2. Direction tuning fits on the MU indicated a weak to strong direction tuning across the channels and shanks. Mean firing rates peaked for direction tunings around 180°, with a median preferred direction of 171.8°.

**Fig. 2.**
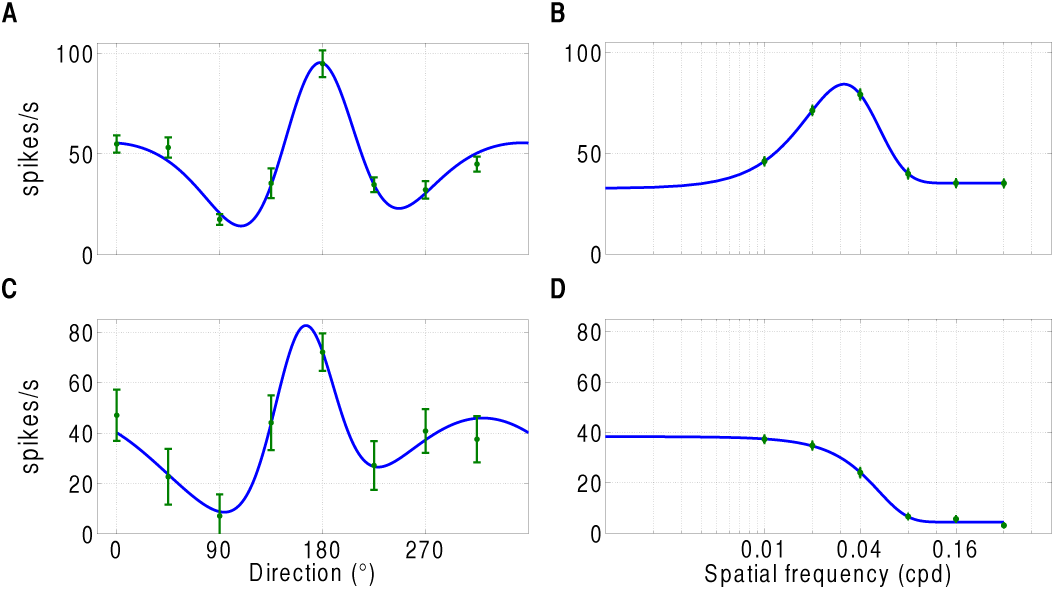
Examples of typical direction tuning curves and their corresponding SF tuning curves on the same site from two mice. (A, C) show strong orientational/directional tuning with a peak around 180^°^. (B, D) reveal the SF tuning on the same electrode sites as in (A) and (C), illustrating strong bandpass and lowpass properties, respectively. Note different y-axes for the two mice. Direction tuning in degrees, SF tuning in units of cpd, error bars standard error of the mean (s.e.m)

In contrast to what has been suggested in the literature [8], the polar plot of preferred spatial frequency against preferred direction indicated that certain directions were overrepresented across the visual field (Fig. 3). Here, the graph of preferred SF against preferred direction of 217 MU for 10 mice revealed a majority of SF/direction pairs appearing clustered around 0.02-0.03 cpd and 180°. The polar plot in (B) of Fig. 3 showing DSI against preferred direction further illustrates this bias towards horizontal moving gratings.

**Fig. 3.**
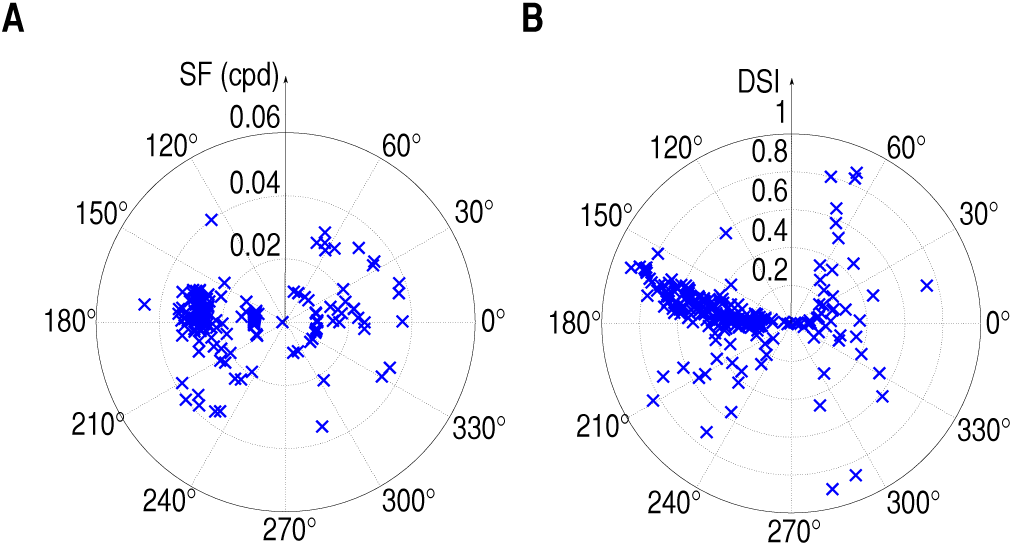
Preferred direction is not uniformly distributed. (A) Polar plot of preferred spatial frequency against preferred direction (217 sites from 10 animals). (B) Polar plot of DSI against preferred direction.

### B. Spatial Frequency Tuning

A subset of MU indicated significant SF tuning. Similar to values reported in the literature [8], the median preferred SF was 0.022 cpd. About half of the MU revealed bandpass properties indicated by a drop in response to the lowest SF, 0.01 cpd, which is illustrated by the example of panel (B) of Fig. 2. Our analysis showed a median cutoff spatial frequency of 0.12 cpd (as determined as the -3 dB cutoff from the preferred SF).

Most MU revealed a preferred SF of around 0.02-0.03 cpd, which is in accordance with results seen in SU studies [1], [8], [12], [13]. Variation was low and only occasionally did we observe a peak SF response exceeding 0.04 cpd, as evident from Fig. 3.

### C. Orientation Selectivity Index

118 MU had an OSI exceeding 0.5. Further, the mean OSI amounted to 0.51 ± 0.01 (s.e.m); the mean DSI to 0.41 ± 0.01 (s.e.m).

### D. Decoding Performance

Decoding performance was determined with a confusion matrix and %-correct. Fig. 4 (A) shows the average confusion matrix for decoding SF from spike responses for the best decoder (Naive Bayes in both cases), where circle size corresponds to %-correct classification. It is visible that low SFs performed better than high SFs, which reflects the low-pass and band-pass properties of the tuning functions. Panel (B) of Fig. 4 reveals the average confusion matrix for directional decoding achieving an evenly high classification performance for all directions. Average performance for SF decoding with the Naive Bayes classifier as depicted in panel (A) of Fig. 4 was 43.7% (chance level 16.67%), whereas the averaged direction decoding performance (B) achieved 58.5% (chance level 12.5%). For the direction decoding task, we classified 80 samples, for which an error rate of <70% against chance level of 12.5% shows a significant classification performance at p<10^−4^. Decoding performance varied across experiments, Fig. 5 reveals averaged decoding performance across animals for both feature types and decoding targets (direction, SF). The dotted line indicates chance level.

**Fig. 4.**
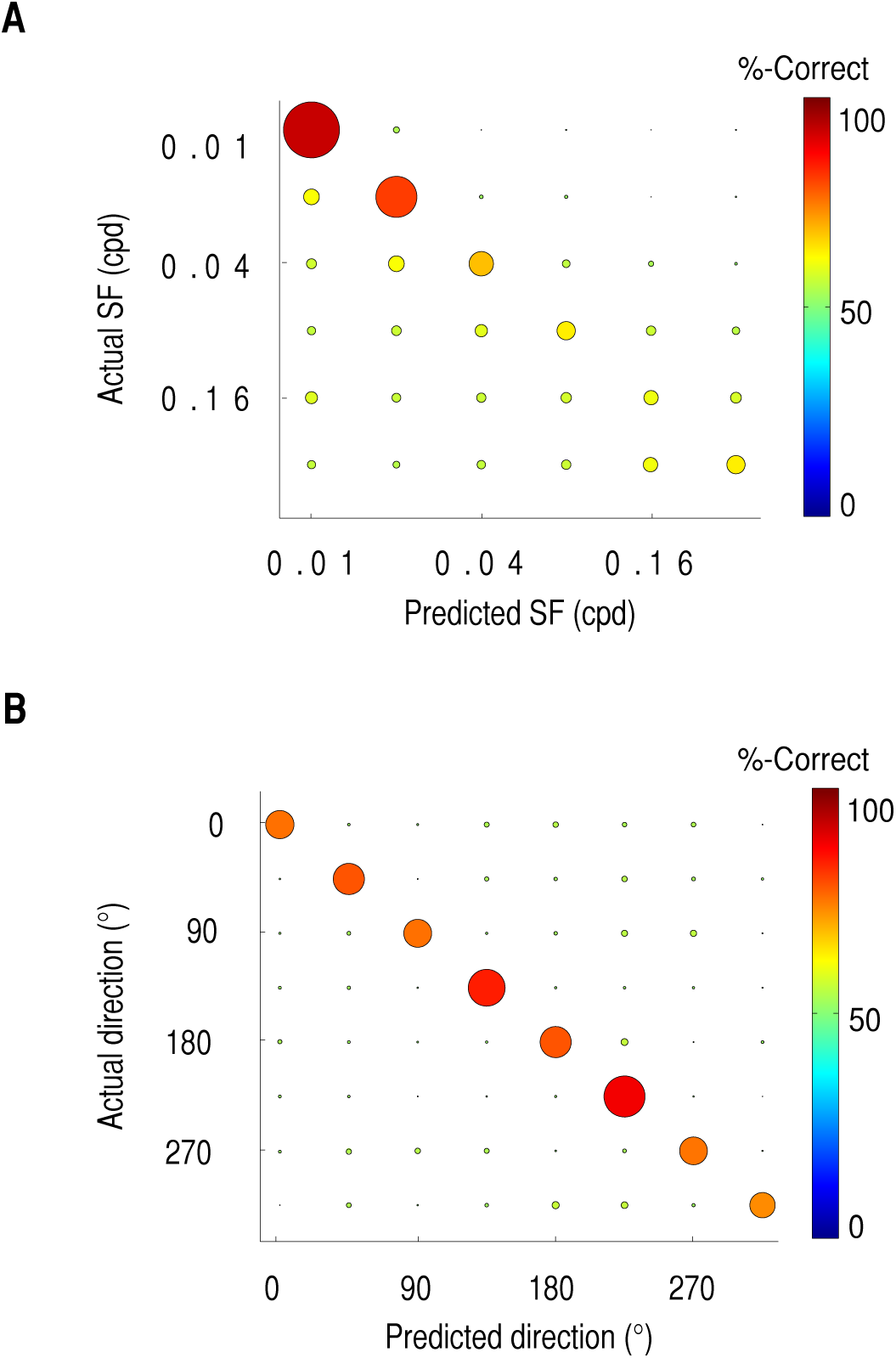
Good predictions for low SFs and high performance on all direction predictions. Confusion matrices for SF (A) and direction decoding (B), where circle radius and colour correspond to correctly classified parameters. (A) shows the averaged confusion matrix of the Naive Bayes classifier on SF decoding, indicating a high prediction success for low SFs, and decreasing success for higher SFs illustrated by smaller circles and colder colours. (B) same as in (A) on direction decoding, demonstrating a consistently high correspondence between predictions and actual directions.

**Fig. 5.**
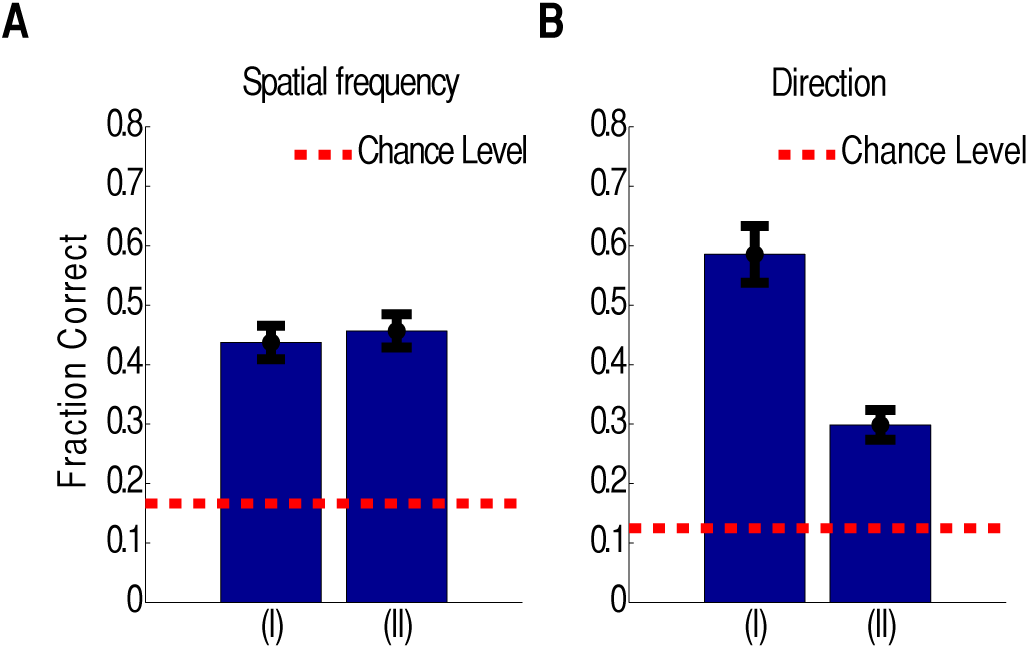
High average classification performance of the best decoder, Naive Bayes, for both features on SF and direction decoding. *A* shows the correct classification rate in SF-decoding from binned spike responses (I), and from the vector of the spike counts of all channels (II). *B*, decoding performance for direction reveals a higher performance for the binned spike response features. Dashed line (red) denotes chance level, error bars in s.e.m.

The different decoders achieved comparable results in decoding SF and direction. Spike train decoding performance varied between 43.7% for Naive Bayes and 34.2% for the kd-tree. The spike count approach for SF decoding ranged between 45.3% and 45.7%. Spike train features achieved between 33.74% and 58.5% correctly detected directions, and spike count feature performance ranged between 26.3% and 38.0%. All results showed statistically significant classification performance (p<0.05).

## V. Conclusions and Discussion

MU recorded from mouse primary visual cortex were selective to spatial frequencies and exhibit bandpass or low-pass properties. With varying modulation depths, preferred direction indicated that specific directions were preferentially distributed around 180° (leftward moving grating) - one result, which is at variance with previous literature [8], [13], [1]. With our MU studies we could confirm the preferred spatial frequency to lie around 0.022 cpd, which has been reported for single units earlier [1], [8], [13]. Our data and analysis of preferred spatial frequency against preferred direction suggests that specific directions were overrepresented across the visual field, particularly around 180°. In addition, a mean OSI of 0.51 and a mean DSI of 0.41 further illustrate the selectivity of MU data. Yet, one may argue that our measure of DSI and OSI may overestimate the true underlying selectivity [14].

We could further demonstrate that MUA can be used to efficiently decode different visual stimuli, with a directional decoding performance of 58.5%. This indicates that MUA contains information about spatial stimulus structure in mouse primary visual cortex. This an important step in utilising information inherent in neural signals for applications, which cannot rely on SU stability, or which do not have the capacities for time consuming spike-sorting. This paves the way for exploring how much information about the stimuli is contained in MU and how they can be efficiently exploited in BMI applications or chronic recordings of behavioural studies, in which it cannot be guaranteed to sustain a stable single unit identification over time.

## Notes

* This work was supported by BBSRC grant BB/K001817/1, and a Royal Society Industry Fellowship to SRS.

